# Integration of lymphatic vasculature to a human lymph node-on-chip enhances physiological immune properties

**DOI:** 10.1101/2025.03.20.644364

**Authors:** Andrew I. Morrison, Jonas Jäger, Charlotte M. de Winde, Henk P. Roest, Luc J.W. van der Laan, Susan Gibbs, Jasper J. Koning, Reina E. Mebius

## Abstract

To study systemic human innate and adaptive immune responses in detail, competent *in vitro* lymph node (LN) models with LN stromal cells (LNSCs) are required to recapitulate the physiological microenvironment. The multicellular organisation of LNs possesses a challenge for designing such microphysiological systems (MPS), particularly with the structural complexity of LNs and the lymphatic vasculature. Here, we established an organotypic LN model with integrated lymphatics in an organ-on-chip (OoC) platform containing a printed sacrificial structure, and studied the influence of a perfused lymphatic endothelial cell (LEC)-lined channel on the LN-on-chip microenvironment. Upon one-week of culture under lymphatic flow, LECs lined the tubular structure forming a lymphatic vessel through the LN model, and stable metabolic conditions within the LN-on-chip were confirmed. Interestingly, LECs in the LN-on-chip displayed the phenotype found in human LNs with upregulation of LEC-specific LN markers, such as atypical chemokine receptor 4 (ACKR4). The presence of the LEC-lined perfused vessel in the LN-on-chip resulted in the increase of native immune cells, most notably B cells, and the secretion of survival and migratory signals, namely interleukin-7 (IL-7) and CC motif chemokine ligand 21 (CCL21). Likewise, LECs promoted the abundance of immune cell clusters closer to the vessel. As such, these features represent an enhanced physiological microenvironment to allow for immune cell migration and interactions for efficient LN functioning. This approach paves the way for LN integration into multi-OoC (MOC) platforms to investigate immunological crosstalk between tissue-derived factors, immune cell trafficking and organ-specific adaptive immune responses.

## Introduction

Lymph nodes (LNs) play an integral part in systemic immunology. Their localisation throughout the human body is tailored towards the efficient drainage of interstitial fluid from all tissues and organs, allowing the screening of pathogens, removal of toxins and the migration of cells to initiate the adaptive immune response. Such immune responses occur due to the highly dynamic and architectural structure of the multicellular LN microenvironment. This organisation is regulated by LN stromal cells (LNSCs) from non-hematopoietic origin^1^. Fibroblastic reticular cells (FRCs), a type of LNSC from the mesenchymal lineage, create the three-dimensional (3D) structural network throughout the LN^2^ and promote crosstalk between dendritic cells (DCs), T cells and B cells for immune activity^3^. In addition to FRCs, the non-mesenchymal LNSCs are composed of endothelial cells (ECs), either blood ECs (BECs) or lymphatic ECs (LECs), with both cell types lining their respective vessels to regulate immune cell migration to and from the LN^1^.

Upon inflammation in peripheral tissues and organs, antigen-induced activation of antigen presenting cells (APCs) can occur, resulting in their migration into the lymphatic vasculature. These are typically DCs, and once activated they upregulate C-C chemokine receptor type 7 (CCR7) to undergo chemotaxis along the afferent lymphatic vasculature into the LN, following a chemotactic gradient of LEC-secreted CC motif chemokine ligand 21 (CCL21)^4^. LECs establish this CCL21 gradient by expressing atypical chemokine receptor 4 (ACKR4) to scavenge CCL21^5^, thereby guiding DCs through the afferent lymphatics and facilitating their arrival into the LN subcapsular sinus (SCS)^6^. Here, DCs enter the LN parenchyma by undergoing transendothelial migration^7^, facilitated by additional chemotactic cues^8^, including CCL19; the second ligand of CCR7^5,9^.

The LECs of the LN are heterogenous and populate intertwined networks of sinus channels across the LN parenchyma. Six subsets of human LECs have been identified based on anatomical location^10^, displaying unique cellular signatures compared to peripheral LECs^11,12^. These include ACKR4^+^ SCS ceiling LECs, TNF receptor superfamily member 9^+^ (TNFRSF9^+^) SCS floor LECs, and macrophage receptor with collagenous structure^+^ (MARCO^+^) medullary LECs. Furthermore, distinct functional properties of LECs have been uncovered in mice, such as producing survival and proliferation signals^13^, trapping and filtering lymph-borne molecules^14^, and exhibiting antigen-presenting capabilities^15,16^. Therefore, LECs of the LN are essential for promoting the adaptive immune response.

To study the LN in health and disease, and recapitulate such an immunological cascade of events for an adaptive immune response *in vitro*, a biological system is required that can reflect the physiological LN environment. Therefore, recent interest has been made in developing LN models to study *in vitro* immunity. Such models are microphysiological systems (MPS), which have been developed in response to the need for better human models for drug testing, disease modelling and basic understanding of organ physiology^17,18^. LN models have currently demonstrated robust humoral immunity from vaccine and cancer therapy applications^19–21^ but are suboptimal based on lack of LNSC inclusion. LNSCs have shown their value in 3D LN models^22–24^, yet these static models lack key physiological elements like flow, which can induce specific cytokine and chemokine expression^25^. In this context, recent advances in the Organ-on-chip (OoC) field become apparent, particularly for using microfluidic devices to model human immunology^26^. LN-on-chips have shown early promise with studying responses to vaccination using peripheral blood mononuclear cells (PBMCs)^27,28^ and tonsil-derived cells^29^, stromal-regulated immune cell migration^30,31^ and cell-cell interactions^32,33^. However, LN-on-chip models that use native LN-derived cells and integrate lymphatic vasculature are currently non-existent.

To resemble the human LN function and architecture more closely, we have previously developed a static LN model with native immune cells and enriched with FRCs^23^. In this study, we take this LN model one step further and incorporated a perfused LEC-lined lymphatic vessel through the LN model using the TissUse’s HUMIMIC Chip2 OoC device, modified with a sacrificial structure^34^. The influence of LECs and flow on the LN microenvironment was characterised over a one-week culture period. In contrast to a LN-on-chip without lymphatics, the LN-on-chip with integrated LECs showed increased lymphocytes, survival and migratory factors, as well as an abundance of immune cell clusters, which are all hallmarks of an optimal LN-like microenvironment. As such, this LN-on-chip demonstrates a degree of immunocompetency previously unobserved in static LN models.

## Material and methods

### Human tissue collection

Human LNs were obtained during liver transplant procedures of cadaveric donors at the Erasmus MC, Rotterdam, The Netherlands, in accordance with the Medisch Ethische Toetsings Commissie (METC) of Erasmus MC (MEC-2014-060). The use of biopsies for research purposes follows the ethical Helsinki Declaration of 1975 standards. All patients (liver transplant recipients) gave written informed consent to use their donor tissue. LNs were resected along the hepatic artery and portal vein in the porta hepatis. LNs were transported in Belzer UW Cold Storage Solution (Bridge to Life Ltd., London, England, UK) and processed within 72 hours of surgery. Donor details are in Table S1.

### Enzymatic digestion of human lymph nodes

Immune cells and LECs were isolated from human LN-tissue by enzymatic digestion, as previously described^35^. Briefly, 4 x 10-minute digestion cycles of LNs in an enzyme mixture containing RPMI-1640 with 2.4 mg/ml Dispase II, 0.6 mg/ml Collagenase P and 0.3 mg/ml DNase I (all from Sigma-Aldrich, St. Louis, MO, USA) was performed. Ice cold phosphate-buffered saline (PBS), supplemented with 2% foetal calf serum (FCS) (HyClone; GE Healthcare, Chicago, IL, USA) and 5 mM ethylenediaminetetraacetic acid (EDTA), was used to stop digestion and wash the isolated cells at 300 G centrifugation for 4 mins at 4 °C. Cell pellet was re-suspended in 1 ml of DMEM (Gibco, Grand Island, NY, USA) with 10% FCS, strained through a 100 µM filter, and counted. The LN cell suspensions were either cryopreserved or cultured as described below.

### Culture of primary lymphatic endothelial cells

LN cell suspensions were seeded at a density of 1.25 x 10^6^ cell suspension per cm^2^ on culture flasks, coated with 2 µg/cm^2^ gelatin (Sigma-Aldrich). For selective LEC outgrowth, culture media comprised of Endothelial Cell Growth Medium MV2 (MV2; PromoCell, Heidelberg, Germany) with 10% FSC and 1% Penicillin/Streptomycin/Glutamine (PSG). After three days, lymphocytes were washed away with PBS, and upon confluence, cells were passaged and harvested with 0.5% Trypsin + 5 mM EDTA. LECs were used up to passage 4 for all individual experiments.

### Microfluidic device

In this study, we made use of the HUMIMIC Chip2 24-well (TissUse, Berlin, Germany) with a printed sacrificial structure (single or bifurcated print) within the larger 14 mm diameter culture compartment (further referred to as the LN compartment) and aligned with the on-chip microcirculation (Figure 1B). The manufacturing details, characterisation of flow parameters and method of endothelialisation have been described previously for this chip^34^. Flow was applied during the experimental culture period in a counterclockwise circulation using a HUMIMIC Starter (TissUse) control unit, set at a 30 bpm frequency with ± 50 mbar to simulate lymphatic flow^34,36^.

**Fig. 1:**
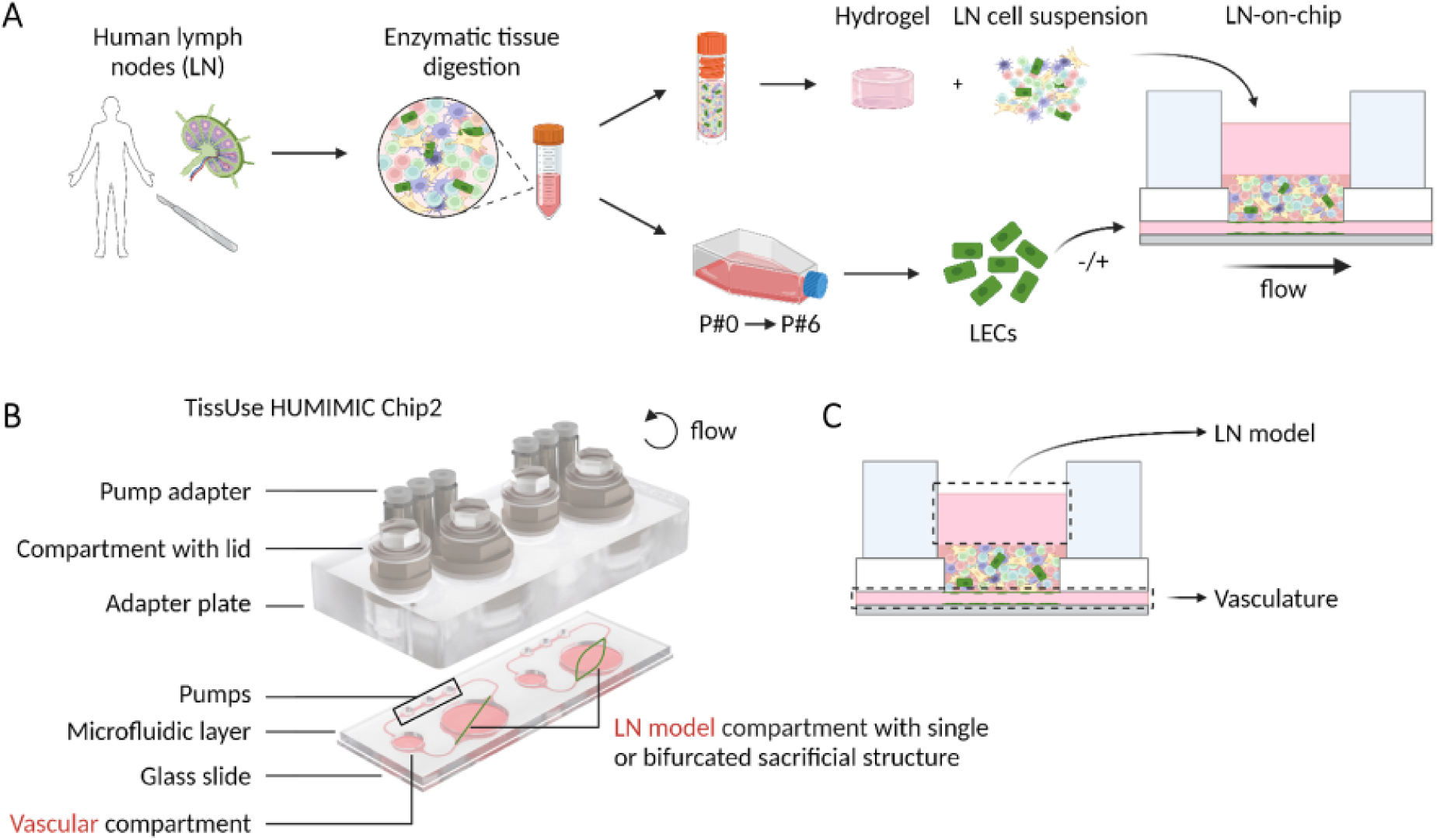
Generation of a vascularised lymph node model on-chip. A. Procedure to isolate cells from a human LN. After enzymatic digestion, the LN cell suspension was cryopreserved and donor-matched LECs amplified until passage 6. Subsequently, the LN cell suspension was thawed, incorporated in a collagen/fibrin hydrogel, and added to a chip platform. Mimicking lymphatics, LECs lined a perfused channel underneath the LN model to generate a vascularised LN-on-chip. LN-on-chip models were compared with either an empty vessel or a vessel lined with LECs. B. Explosion view of the HUMIMIC Chip2 chip platform used in this study. In the LN compartment, a sacrificial structure was incorporated to create a lymphatic channel containing LECs underneath the LN hydrogel. C. Sampling in the vascular compartment and the LN compartment allows to separate circulating medium in the vasculature and static medium on top of the LN model. Created with BioRender.com

### Construction of LN-on-chip model

The 3D LN model was constructed, as previously described^23^, directly into the HUMIMIC Chip2’s LN compartment (Figure 1A). This was performed by mixing human LN cell suspensions with a hydrogel composed of rat-tail collagen type I (3 mg/ml end concentration) in Hank’s Balanced Salt Solution (HBSS without Ca and Mg, Gibco) and fibrinogen from human plasma (1 mg/ml end concentration; Enzyme research laboratories, cat no: FIB 1, South Bend, IN, USA). LN cell suspensions were adjusted to a concentration of 10 x 10^6^ cells per ml of hydrogel, with a final hydrogel volume at either 300 or 350 µl. Thrombin (0.5 U/ml; Merck KGaA, Darmstadt, Germany) and aprotinin (50 KIU/ml; SERVA, Heidelberg, Germany) were added to the hydrogel for fibrinogen polymerisation and to prevent shrinkage respectively. Hydrogels were polymerised inside chips at 37 °C and 5% CO_2_ for 30 mins, after which the sacrificial structure dissolved to leave a hollow channel. Complete MV2 medium was added to the vascular compartment (300 µl) and LN model medium, composed of RPMI-1640 with HEPES and L-glutamine (Gibco), 10% FCS (HyClone; GE Healthcare), 2% PSG, 1% ITS (Gibco), 1X non-essential amino acids (NEAA) (Gibco), 1X sodium pyruvate (Gibco) and 1X normocin (Invivogen, Toulouse, France), was added to the hydrogel compartment (500 µl). Both media contained 50 KIU/ml aprotinin (to reduce shrinkage). The chip was connected to the HUMIMIC Starter for an overnight incubation at 37 °C and 5% CO_2_ in flow (± 500 mbar, 30 bpm) to remove remnants of the dissolved sacrificial structure in the lumen.

### Lymphatic endothelial cell seeding

The next day, remaining residuals of the printed sacrificial structure were removed by a medium exchange of MV2 in the vascular compartment. LECs were then seeded at a total number of 5 x 10^5^ cells to the vascular compartment and flushed through the vessel under the LN model via pressure differences applied with the compartment lids. Chips were incubated at 37 °C, 5% CO2 for 4 hours. After incubation, the vascular compartment was washed with MV2 medium. Next, 300 µl of fresh MV2 medium was added to the vascular compartment, and 500 µl of fresh LN model medium on top of the hydrogel, with 50 KIU/ml aprotinin (to reduce shrinkage) supplemented to both media from here onwards. LN-on-chip models without LEC endothelialisation followed the same procedure.

Chips were then cultured under flow at 30 bpm ± 50 mbar for a seven-day period. A full medium exchange was performed on day 3 and 5 for both compartments, with each compartment supernatant stored separately at 4 °C and −20 °C for future analysis (Figure 1C). At the end of the culture period, medium from the two compartments was collected separately and the LN-on-chip was either fixed *in situ* for 3D imaging, or the LN hydrogel was removed for enzymatic digestion. Enzymatic digestion to a single cell suspension for analysis by flow cytometry was achieved using 0.2 mg/ml Collagenase P and 0.1 mg/ml DNase I for 1 hour at 37°C, after which the reaction was stopped with PBS containing 2% FCS and 5 mM EDTA.

### Metabolic analysis

Levels of glucose, lactate, and lactate dehydrogenase (LDH) activity were analysed from chip supernatants on day 3, 5 and 7. This was performed using a Glucose Colorimetric Detection Kit (EIAGLUC, Thermo Fisher), Lactate Assay Kit (MAK064, Sigma-Aldrich) and Cytotoxicity Detection Kit PLUS (LDH) (04744934001, Roche, Basel, Switzerland), according to the manufacturer’s instructions. MV2 and LN model medium were used as background controls in each analysis.

### Flow Cytometry

Cell suspensions were stained in a 96-well U bottom plate at 4 °C for flow cytometric analysis. Cells were first washed with FACS buffer containing PBS, 0.1% bovine serum albumin (BSA) and 0.05% NaN_3_, and stained with a fixable viability dye (eFluor™ 780; Invitrogen; #65-0865-14) for 10 mins at 4°C. Fc-receptor blocking was performed using 10% normal human serum, and cells were then incubated with directly labelled antibodies. Post-staining, cells were washed two times with FACS buffer and fixed with 2% paraformaldehyde (PFA) (VWR, Radnor, PA, USA) for 10 mins at room temperature (RT). An overview of all antibodies used can be found in Table 1. Samples and single stains on beads were acquired on Aurora 5-laser Flow Cytometer (Cytek; Fremont, CA, USA). Autofluorescence (AF) correction of cells was performed as previously described^37^. Data analysis was conducted using OMIQ (Boston, MA, USA).

**Table 1.**
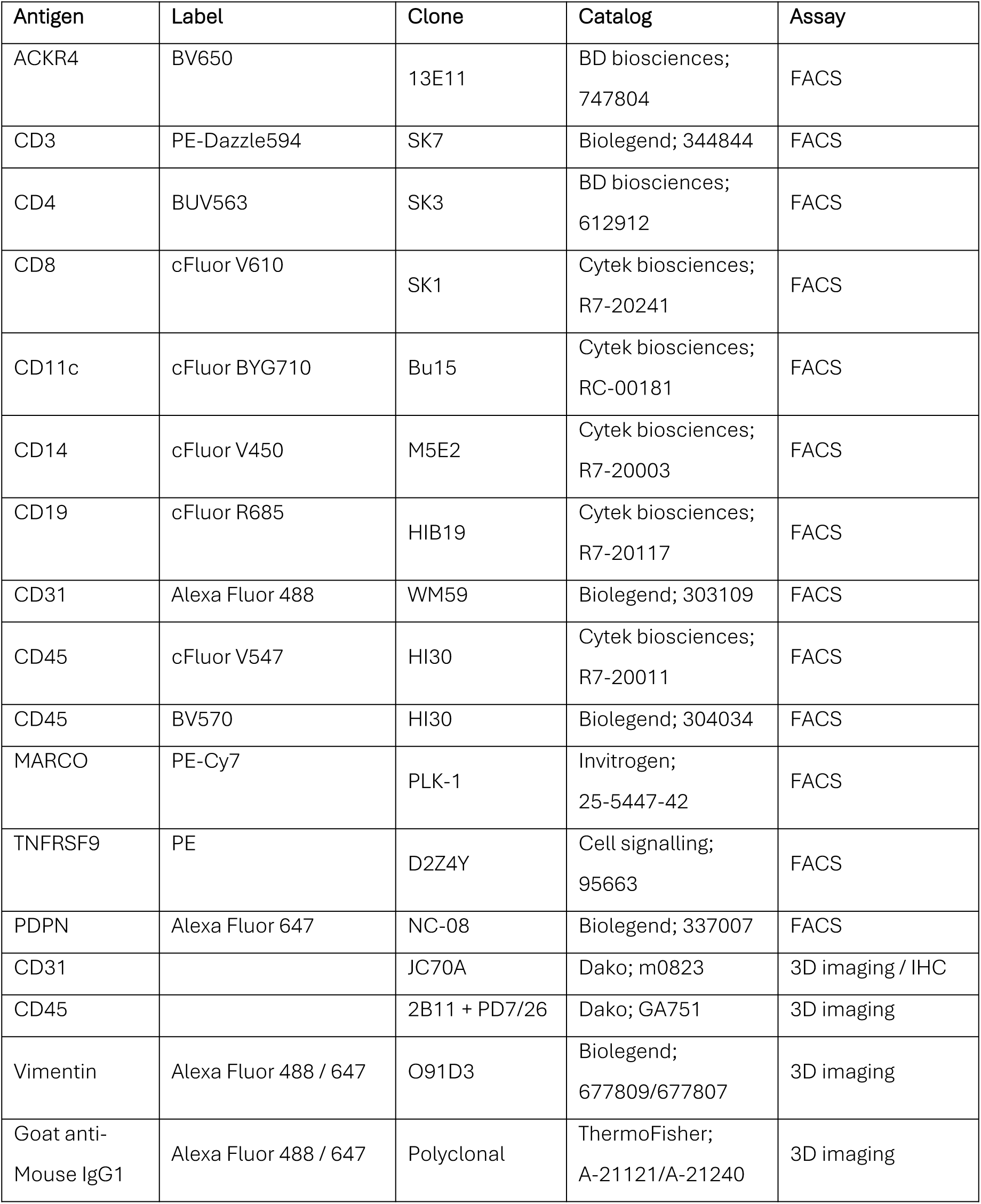
Overview of antibodies. FACS: Flow cytometry, IHC: Immunohistochemistry.

### Immuno-fluorescent and -histochemical staining

The LN-on-chip hydrogel and channels were fixed in the chips by directly adding 4% PFA in PBS (Electron Microscopy Sciences, Hatfield, PA, USA) in PBS for 10 min under flow (± 500 mbar, 30 bpm) followed by 20 mins static at 37°C. Chips were then washed twice with PBS for 10 mins pumping at RT (± 500 mbar, 30 bpm). Hydrogels were then preserved in PBS with 0.02% NaN_3_ before antibody staining or paraffin embedding. An overview of all antibodies used can be found in Table 1.

For *in situ* immunofluorescent chip staining, chips were pre-treated with 0.2% Triton in PBS, followed by incubation in the chip with an unconjugated antibody to both the vascular (100 μl) and LN (200 μl) compartment, diluted in 0.2% Triton in PBS. Unconjugated antibodies used were anti-CD31 (clone JC70A, Dako; m0823) and anti-CD45 (clone 2B11 + PD7/26, Dako; GA751). Staining was performed under flow (± 500 mbar, 30 bpm) for 30 mins at RT to ensure complete antibody coverage throughout the microfluidic circuit, then left overnight at 4 °C on a rocker to allow antibody diffusion through the hydrogel. The next day, chips were washed twice with 0.2% Triton in PBS and incubated with a secondary antibody conjugate at RT on a rocker for 2 hours. The secondary antibody conjugate used was goat anti-mouse IgG1 (Alexa Fluor 488/647, polyclonal, ThermoFisher; A-21121/A-21240). Afterwards, a directly labelled antibody, anti-Vimentin (Alexa Fluor 488/647, clone O91D3, Biolegend; 677809/677807) was added to the chip for an overnight incubation at 4 °C on a rocker. Before image acquisition, DAPI (Invitrogen) was added for 1 hour at RT, followed by two washing steps and storage at 4 °C until analysis. Chips were imaged by confocal microscopy using a Nikon AXR (Nikon, Tokyo, Japan).

For paraffin embedding, hydrogels were removed from the chip and subsequently dehydrated, embedded in paraffin and cut at 5 μm sections for immuno-histochemical (IHC) analysis. For CD31 staining, sections were deparaffinised and immersed in 10 mM Tris/1 mM EDTA buffer (pH 9.0) for 15 mins at 100°C for antigen retrieval, followed by slowly cooling to RT. Sections were then washed with PBS and stained for 1 hour with anti-CD31 (clone JC70A, Dako; m0823). This was followed by incubation with BrightVision plus Poly-HRP-Anti-Mouse/Rabbit/Rat IgG (Immunologic, VWR International B.V., Breda, the Netherlands) and 3-amino-9-ethylcarbazole (AEC, Sigma-Aldrich) substrate, concluded with a hematoxylin counterstain. Sections were imaged using a Nikon Eclipse 80i microscope (Nikon, Tokyo, Japan) with NIS Elements 4.13 software (Nikon).

### Image analysis

Immunofluorescent 3D images were analysed using Imaris (v10.1.0; Oxford Instruments, Oxfordshire, UK). Firstly, the vessel surface was rendered with machine learning segmentation (10 µm smoothness detail) based on DAPI for models without a LEC vessel, and Vimentin for models with a LEC vessel. Subsequently, spots were generated on DAPI^+^CD45^+^ cells to identify immune cells (4.5 µm approximate diameter). A cell cluster was defined as one cell with nine cell neighbours within an average distance of 20 µm between each cell. Then, the average distance of immune cells or clusters to the vessel surface was calculated.

### Cytokine/Chemokine analysis

Chip supernatants were collected separately from the vasculature and hydrogel compartment and analysed for the presence of cytokines and chemokines. This was performed using either a cytokine bead array (CBA; Custom LEGENDplex panel, BioLegend) or an enzyme-linked immunosorbent assay (ELISA) for CCL21 (#ab193759, Abcam, Cambridge, UK), according to the manufacturer’s instructions. CBA acquisition was performed on AttuneNxT (Thermofisher, Waltham, MA, USA) and protein concentrations were determined using the LEGENDplex Data Analysis Software Suite (BioLegend).

### Statistical analysis

All data are presented as mean ± standard error of the mean (SEM). The number of human LN donors is described in each figure legend. Statistical analysis was conducted using GraphPad Prism 9 software version 9.5.1 (GraphPad Software Inc., La Jolla, CA). Statistical tests are indicated in figure legends. Differences were significant when P < 0.05.

## Results

### Development of metabolically stable LN-on-chip with integrated lymphatics

A schematic construction of the LN-on-chip with integrated lymphatics is illustrated in Figure 1. Briefly, the LN hydrogel model^23^ was first cast into the LN compartment of the HUMIMIC Chip2, after which the hollow channel of the sacrificial printed structure was seeded either with or without donor-matched LECs. These two biological conditions allowed for studying the influence of LECs on the LN-on-chip microenvironment over seven days under flow. Conditioned media samples were collected from both the circulating lymphatic vasculature and above the LN hydrogel.

To determine whether the LN-on-chip was viable and metabolically active, LDH, glucose and lactate concentrations were measured in the supernatant from both compartments at day three, five and seven of culture. A higher concentration of LDH was measured in both chip compartments at day three in the LN-on-chips with a LEC vessel, which then decreased over the seven days in the vasculature (Figure 2A and S1A). Glucose consumption from the initial cell culture medium remained similar for both conditions, with no clear changes over time (Figure 2B and S1B). More lactate was secreted in the models with a LEC vessel compared to without, which was higher in the vasculature compartment compared to the LN model (Figure 2C and S1C).

**Fig. 2:**
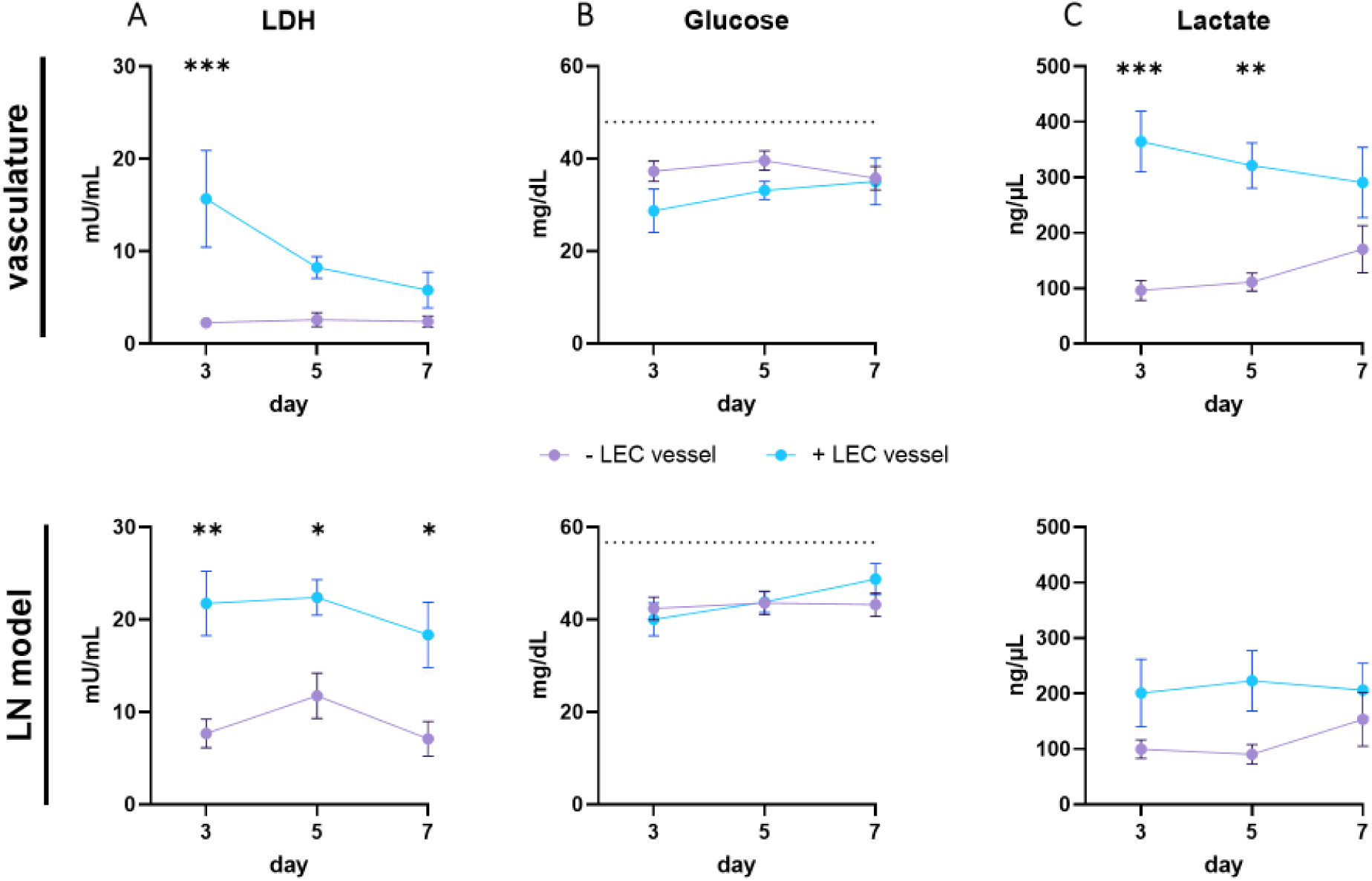
LN-on-chip is viable and metabolically stable for 7 days with and without a LEC vessel. A. Lactate dehydrogenase (LDH), B. Glucose (with initial levels from culture media indicated as dotted line), C. Lactate concentrations in the culture medium, measured at day 3, 5 and 7 during the LN-on-chip culture period. Medium was sampled either from the perfused vasculature (upper graphs) or on top of the LN model (lower graphs). Shapes represent different donors as mean ± SEM; n = 4 independent experiments, each performed with intra-experimental duplicates. Significance indicated between ± LEC vessel per day with ordinary two-way ANOVA (*\*p* < 0.05, *\*\*p* < 0.01, *\*\*\*p* < 0.001).

As such, these factors collectively showed that, within a one-week culture period under flow, a stable metabolic profile of a LN-on-chip with an integrated LEC vessel was generated.

### LECs in LN-on-chip physiologically represent LN lymphatics

Next, to determine whether the unique properties of LN LECs were recapitulated in the LN model, morphology and phenotype of the LECs were characterised within the LN-on-chip. Upon confocal imaging of the endothelialised LN models, a uniform adherence of LECs along the inner layer of the vessel wall and a visibly distinct lumen structure was observed from CD31 and Vimentin staining (Figure 3A and 3B). These LN-derived LECs have also been previously described to express VE-cadherin^+^ tight junctions and permeable integrity when in a similar vessel-like setting^38^. The lumen structure remained present in vessels without LECs (Figure 3B and S2A), indicating vessel stability through the LN model regardless of LEC endothelialisation. The vessel width was slightly larger with LECs (453 µm ± 76 µm) compared to without LECs (348 µm ± 26 µm) (Figure 3C), with both mean measurements deviating above and below the initial 420 µm vessel width reported after printing^34^, respectively.

**Fig. 3:**
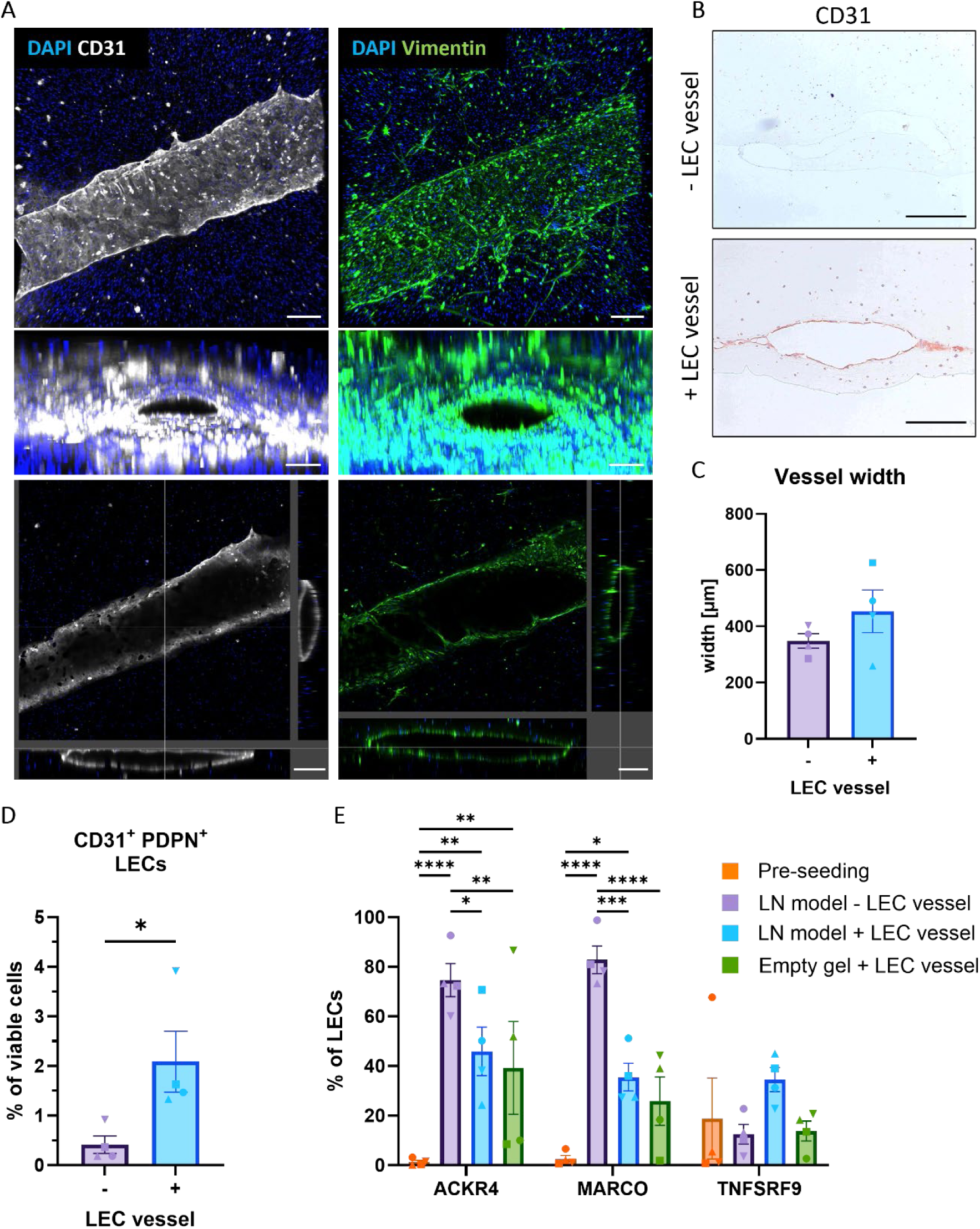
Lymphatic endothelial cells physiologically represent LN lymphatics on-chip. A. LEC vessel staining for DAPI, Vimentin and CD31. Depicted are z-stacks with a top view and a cross section of the channel and a section view, indicating xy-, xz- and yz-cross sections. Scale bars: 200 µm. B. Immunohistochemical staining for CD31, images −/+ LEC vessel show cross-sectional area of channel. Scale bars: 100 µm. C. Width of the channel in the LN-on-chip model without or with (−/+) a LEC vessel. D. Proportion of LECs in the LN-on-chip model −/+ LEC vessel. E. Detailed characterisation for SCS ceiling (ACKR4), medullary (MARCO) and SCS floor (TNFSRF9) LEC markers before seeding in the chip, and after a seven-day culture period on-chip with LN model −/+ LEC vessel or + LEC vessel with an empty hydrogel. Representative gating strategy and histogram overlays for these three markers found in (Figure S2C and S2D). Shapes represent different donors and columns mean ± SEM; n = 4 independent experiments. Ordinary two-way ANOVA (*\*p* < 0.05, *\*\*p* < 0.01, *\*\*\*p* < 0.001, *\*\*\*\*p* < 0.0001).

The phenotype of the LECs from the LN-on-chip model was further investigated after digestion of the LN hydrogel to a single cell suspension using flow cytometry. Prior to LEC-endothelisation of LN-on-chip model, an average purity of 95% LECs across donors from *ex vivo* cultures was confirmed based on CD31 and podoplanin (PDPN) expression (Figure S2B). Within the LN-on-chip hydrogel model with an integrated LEC vessel, CD31⁺PDPN⁺ LECs were identified as constituting to approximately 2% of the total viable cells (Figure 3D). In the absence of a LEC vessel, 0.5% of total cells in the hydrogel were LECs, which is the physiological-like proportion within the native LN cell suspension^35^.

To further characterise the LECs in the LN-on-chip model, we analysed the expression of specific markers associated with distinct LEC subsets found in human LNs, i.e. MARCO, ACKR4 and TNFSRF9^10^. Interestingly, compared to the LEC phenotype prior to endothelisation from *ex vivo* cultures, significantly more LECs expressed ACKR4 and MARCO observed in both LN-on-chip models with and without a LEC vessel (Figure 3E). It is worth noting that the small number of LECs in the total cell suspension from the LN model without a LEC vessel (0.5% in Figure 3D) had the highest expression of both markers. To investigate whether expression of these LEC markers was due to flow and/or presence of immune cells within the LN model, an empty hydrogel with a LEC vessel was generated to simulate a flow-only condition. In the absence of the LN cell suspension, most LECs lost their CD31^+^PDPN^+^ phenotype (Figure S2B), but those remaining as LECs still expressed higher ACKR4 compared to cultured LECs (Figure 3E). No changes to TNFSRF9 expression were seen on LECs across all conditions.

In summary, these results highlight an intact LEC vessel in a LN model and demonstrate a more physiological LEC phenotype on-chip, with characteristics resembling SCS ceiling and medullary LECs.

### LECs preserve B cells within LN-on-chip

To determine the composition of the native immune cells within the LN-on-chip, and whether this was influenced with the integration of a LEC vessel, general immune cell populations were phenotyped by flow cytometry analysis (Figure S2C). Most immune cells in the LN-on-chip models were found to be T- and B-lymphocytes, with a degree of donor heterogeneity (Figure 4A). T cells were comprised of more CD4^+^ than CD8^+^ T cells (Figure 4B), and the percentage of myeloid cells was 1% and 0.5% (−/+ vessel, respectively) of the overall immune cells. While the presence of the LEC vessel did not significantly alter the composition of immune cells, it did significantly increase the total number of immune cells after the seven-day culture (Figure 4C). This was most striking for the B cells, evident from the respective fold change increase compared to the same donors without a LEC vessel (Figure 4C).

**Fig. 4:**
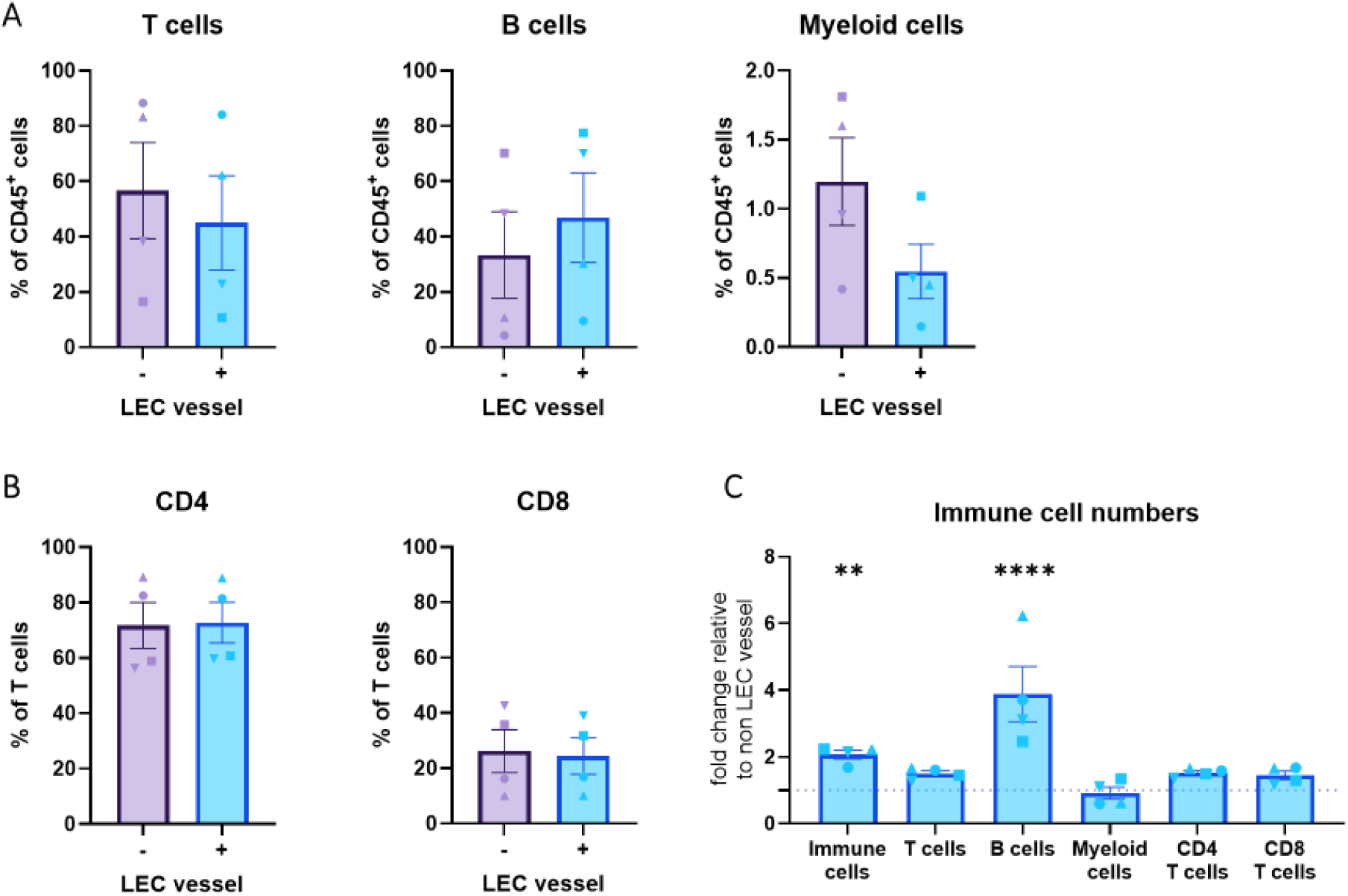
Consistent immune cell populations in LN-on-chip, independent of LEC vessel presence. A. Populations of CD3^+^ T cells, CD20^+^ B cells and CD11c^+^CD14^+^ myeloid cells within the CD45^+^ immune cells after a 7 day culture on-chip −/+ LEC vessel. B. Composition % of CD4 and CD8 T cells −/+ LEC vessel. C. Fold change in number of immune cells in LN-on-chip model + LEC vessel, relative to - LEC vessel set at 1. Shapes represent different donors and columns mean ± SEM; n = 4 independent experiments. Ordinary two-way ANOVA (*\*\*p* < 0.01, *\*\*\*\*p* < 0.0001).

Henceforth, this shows that immune cell populations, and especially B cells, increase in number with the existence of a LEC-populated lymphatic vessel, indicating a successful generation of an immunocompetent LN-on-chip model.

### LECs induce LN-relevant homeostatic factors within LN-on-chip

Since the ability of immune cell function in the LN requires LEC-secreted survival and migratory signals^7^, cytokine and chemokine analysis of the LN-on-chip with and without a LEC vessel was performed at day three, five and seven from both the vasculature and the LN model compartment. LN-on-chips with a LEC vessel revealed an increased secretion of the survival cytokine IL-7 in both compartments on day three and five in the LN model (Figure 5A). The total cumulative secretion of IL-7 throughout the system across one-week was significantly higher in the LN model with a LEC vessel (Figure 5B). Higher concentrations of CCL21 were found in the vasculature at day three within LN-on-chip with a LEC vessel but decreased over time. Total CCL21 secretion was higher in the vasculature with a LEC vessel than no LEC vessel. Interestingly, homeostatic LN-relevant factor CXC motif chemokine ligand 12 (CXCL12) was highest on day five in the LN model with a LEC vessel, and total CXCL12 secretion was significantly more pronounced in the LN model with a LEC vessel compared to the vasculature and the LN model compartment without a LEC vessel. The B cell chemoattractant CXCL13 was found at a higher concentration in the LN model compartment compared to the vasculature, but there was no direct LEC-vessel influence.

**Fig. 5:**
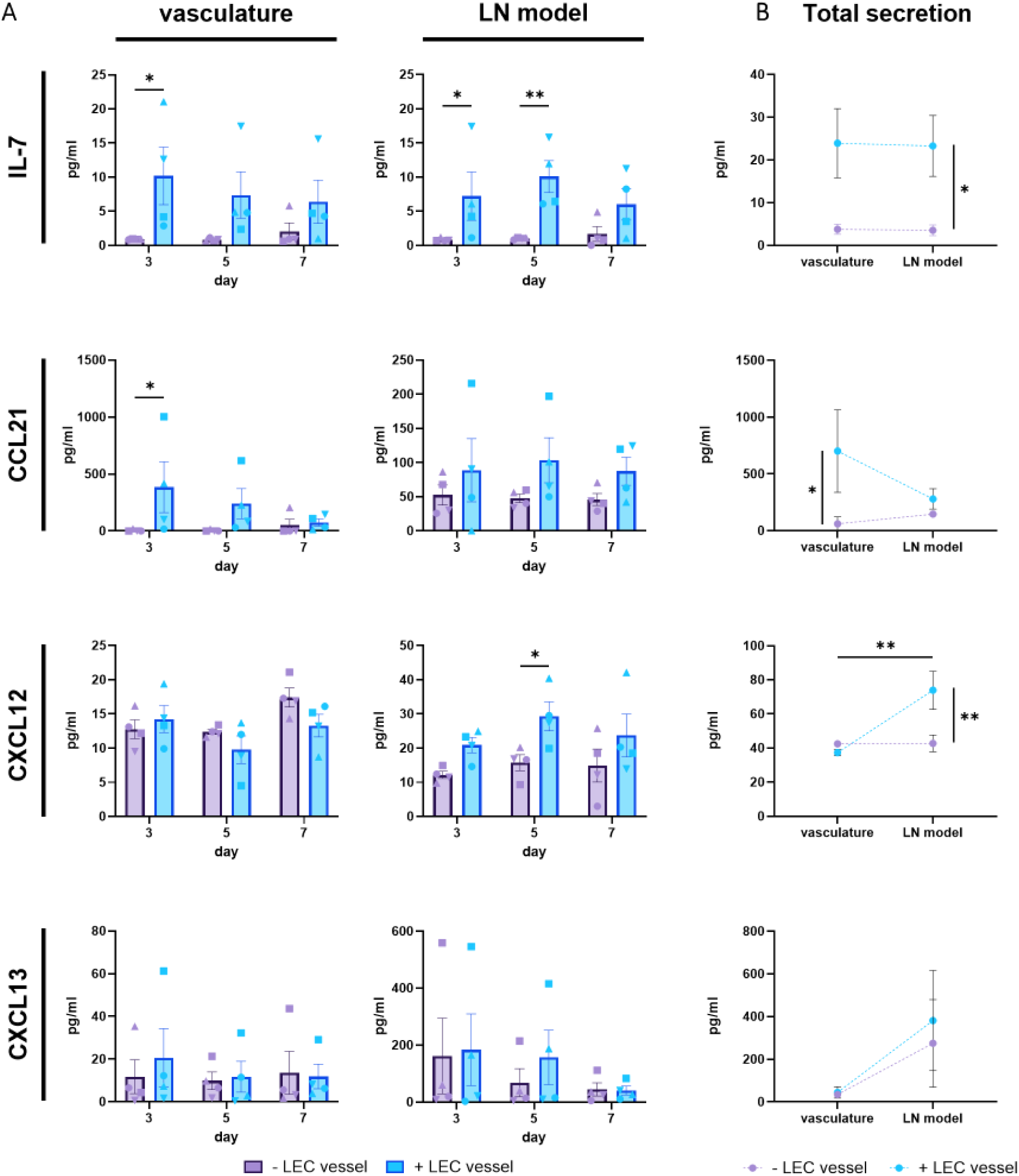
Addition of a LEC vessel results in increased IL-7 and CCL21 secretion in the vasculature. A. LN-specific cytokine secretion, IL-7, CCL21, CXCL12 and CXCL13, into the vasculature (left graphs) and hydrogel (right graphs) supernatant of the chip throughout the culture period at day 3, 5 and 7. Compared are empty hydrogel channels with a lymphatic vessel. B. Total cumulative cytokine secretion throughout the 7-day LN-on-chip culture period. Shapes represent different donors and columns mean ± SEM; n = 4 independent experiments of pooled intra-experimental duplicates. Significance of total secretion indicated between ± LEC vessel per compartment (vertical line) or between compartments + LEC vessel (horizontal line), with ordinary two-way ANOVA (*\*p* < 0.05, *\*\*p* < 0.01).

This indicates that an integrated LEC vessel not only supports the production of survival and migratory signals in a LN-on-chip, but also the establishes compartmentalised microenvironments of lymphatics and LN, related to specific LN physiology.

### LECs promote closer proximities of immune cell and cluster formation in LN-on-chip

LNs are structured to promote immune cell interactions, which is essential for optimal immune responses^39,40^. Therefore, the spatial distribution of immune cells in our LN-on-chip models around the lymphatic vessel was investigated. Images were acquired around the middle of the LN model, with a given field of view accounting for a fraction of the whole LN compartment. Here, many CD45^+^ immune cells were found distributed throughout the model (Figure 6A), irrespective of a present LEC vessel (Figure S3A), with some immune cells observed in close vicinity to the lymphatic vessel in certain areas. Quantification revealed that the total number of immune cells in the whole field of view was indifferent between −/+ LEC vessel, yet the presence of a LEC vessel increased the total number of immune cells within < 200 µm proximity of the lymphatic vessel in all donors but one (Figure 6B). Immune cells found grouped together were further defined as an immune cell cluster, with the criteria for one immune cell to have nine neighbours within an average cell distance of 20 µm, that includes overlapping neighbours (Figure 6C, Figure S3B-D). The LN-on-chips with a LEC vessel provoked a higher abundance of immune cells clusters in all donors, and resulted in shorter distances of these clusters to the vessel surface in three out of four donors (Figure 6D).

**Fig. 6:**
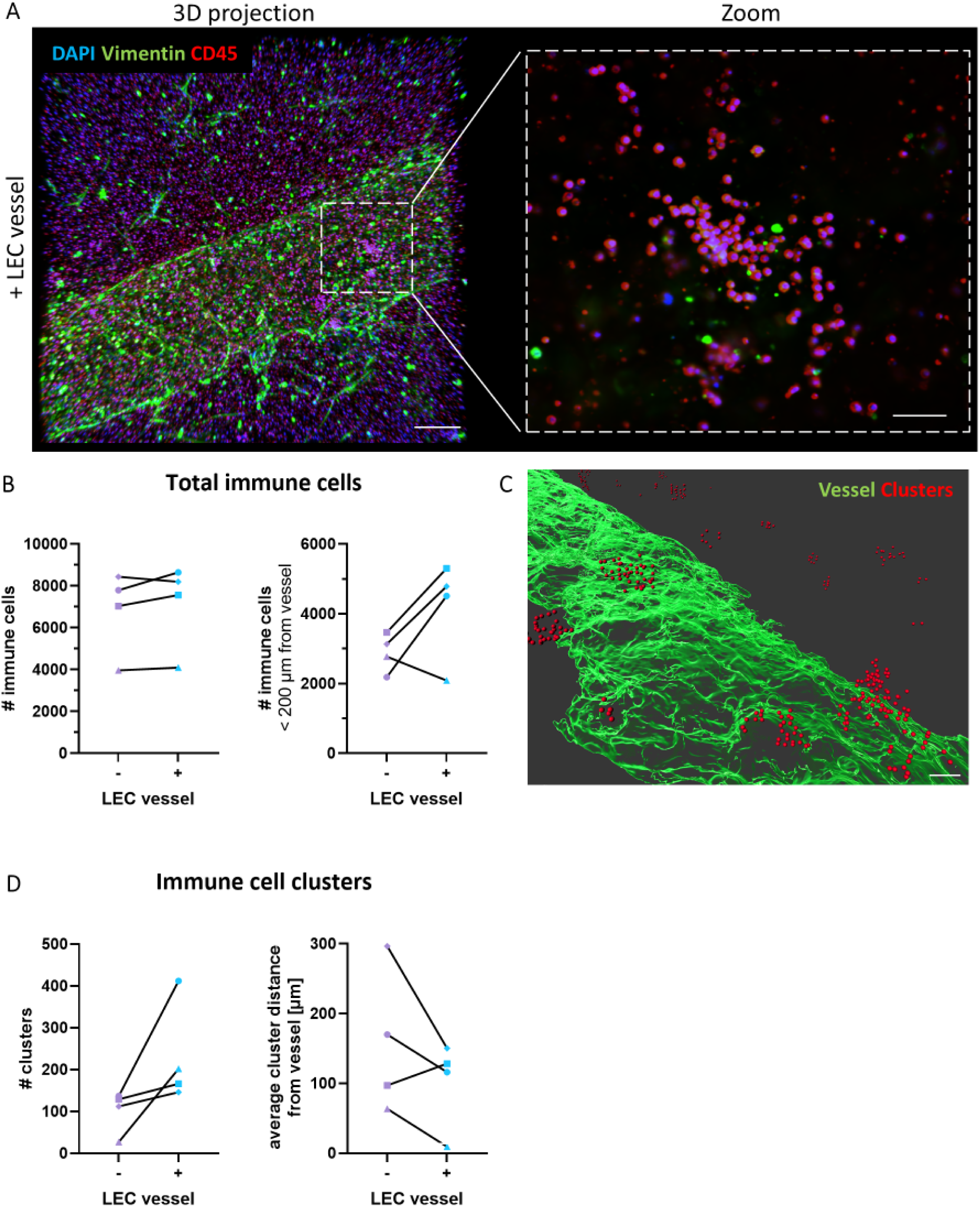
Vascularisation with LECs promotes immune cell clustering. A Staining of the LN-on-chip at day 7 for CD45 (red), vimentin (green) and DAPI (blue) shown as 3D projections + LEC vessel. Zoom magnification visualises immune cell clusters in a single z-plane. Scale bars: 200 µm in overviews, 50 µm in magnification. B. Quantification of the total immune cells and within 200 µm of the vessel. C. Surface rendering for the LEC vessel (green) and clustered immune cells (red) in close proximity of the vessel. Scale bar: 70 µm. D. Quantified clustered immune cells and their shortest distance to the vessel surface. Details are shown in Figure S3.

Therefore, the addition of a LEC vessel seemed to benefit the organisation of the immune cells in proximity to each other and towards the lymphatics, thus in position for optimal immune activity.

## Discussion

The *in vitro* study of systemic immunology requires physiologically relevant human LN models that contain LNSCs and recapitulate organ characteristics^26^. Here we used a commercially available chip system, the HUMIMIC Chip2 24-well with a sacrificial structure from TissUse, to generate a LN-on-chip model with integrated lymphatics. This was achieved by using donor-matched LECs to endothelialise a hollow channel through a LN hydrogel containing a native LN cell suspension. A one-week culture under flow displayed a stable metabolic profile, and LECs added to the LN-on-chip exhibited morphological and phenotypical features that were representative of LN LECs. In addition, LN-on-chip models with a LEC vessel provided the necessary components for LN functioning and immune cell interactions, such as compartmentalised migratory signals, survival factors and immune cell clusters in closer proximity to the lymphatic vessel.

The integration of vascularised organ models is currently a major target within the MPS field^41^. This allows for the delivery of nutrients, homeostatic factors, metabolites and removal of waste products, as well as mimicking processes such as cell trafficking, pharmacodynamics and tissue crosstalk^42,43^. Likewise, it can be used to investigate disease progression, such as tumour metastasis^44^ and EC biology under flow^36,45,46^. Modelling vascularisation has been made possible through various microfabrication techniques, 3D-bioprinting, sacrificial moulds or by inducing cell self-organisation^47^. Regarding the function of the LN, numerous waterfall-like and blind-ended lymphatic channels are intricated through the LN parenchyma to filter the incoming lymph^7,48^, which makes it difficult to accurately model LN lymphatics *in vitro*^49^. This complexity has hindered the development of immunocompetent human LN models since currently they do not address the inclusion of primary LECs and vasculature integration. Here, we began with mimicking lymphatic vasculature in a LN-on-chip model with a LEC-populated channel. As such, even without applying any stimuli to induce immune activity, our data provides evidence that inclusion of a LEC-lined vessel in the LN-on-chip model has a substantial effect on the LN microenvironment during homeostasis.

The organotypic LN model used in this study was comprised of a LN-derived cell suspension which contained both native immune and non-immune cells, such as FRCs and LECs^35^. In contrast to other studies which use PBMCs^27,28^, tonsils^29,33^ or cell lines^30–32,38^ to model a LN-on-chip, we incorporated the cell suspension of native LNs. The static culture of the organotypic LN model was characterised previously by our group^23^, which revealed the benefit of further enrichment with autologous FRCs for viability of lymphocyte subsets, and presence of survival and homeostatic factors. For this LN-on-chip study, the LN model was not further enriched with extra FRCs due to their contractile properties in hydrogel environments^50^. This decision aimed to prevent remodelling or disruption of the lymphatic vessel during the culture period, thus also limiting the culture length to one week. Henceforth, future studies should be aimed at including cultured FRCs and preventing shrinkage through use of other ECM-scaffolds^24,51,52^. Nevertheless, even without FRC enrichment, native immune cells were still observed to be present and maintained in a viable LN environment for the entire culture period, supported by the presence of LECs.

The presence of the lymphatic vessel proved valuable for lymphocytes, particularly for B cells since more cells were found in the hydrogel of the LN-on-chip. The high concentrations of B cell-related factors CXCL12 and CXCL13 most likely account for the maintenance of B cells, which were also originally found in our static LN models along with IL-6, a key B cell survival cytokine^23^. In parallel, the detection of IL-7 in the LEC-populated LN-on-chips was an interesting finding given its previous absence in our static LN models^23^. Both IL-7 and CCL21 are produced by LECs, but while CCL21 is highly enriched within the vasculature, the levels of IL-7 are comparable in both compartments. While IL-7 is most commonly known for naïve T cell survival^53,54^, it can also influence B cells during LN organogenesis^55^, as well as their development^56^ and maturation^57^. The addition of LECs resulted in more CCL21 secreted into the system, with low levels of CCL21 found in the LN model which was also previously absent in static cultures. Flow has been reported to induce CCL21 expression in cultured FRCs^25^, yet it remains to be determined here if LEC secretion of CCL21 is enhanced by flow. However, this would require testing in a different experimental set-up, as it is not feasible to maintain the LN-on-chip under static conditions due to rapid stagnation of culture medium, which reduces cell viability. Both IL-7 and CCL21 are crucial for optimal LN functioning *in vivo*, as knockout mouse models of these factors demonstrated severe depletion of lymphocytes and immune malfunctioning under antigen challenge^58–60^. Furthermore, other LN-on-chip models lacking LECs failed to capture presence of IL-7 and CCL21 in the system^28,30,32^. Therefore, the emergence of these factors associated with the normal physiology of functional LNs, after the integration of a LEC vessel into a LN-on-chip, indicates an improved *in vitro* LN model to generate a supportive microenvironment for lymphocyte survival and potential migration ability.

Additionally, LECs promoted the clustering of immune cells in close proximity to the lymphatic vessel in the LN-on-chip, another physiological attribute for LN functioning^39,40^. In the native LN, naïve lymphocytes routinely survey soluble antigens or incoming APCs from afferent lymphatics. This requires numerous immune cell interactions for cognate antigen matching, so therefore the increased abundance of immune clusters facilitates such interactions and thus antigen-specific recognition by lymphocytes close to the lymphatic vessel. This is particularly relevant for *in vitro* model functionality in future experiments, as cell density can enhance the likelihood of antigen-specific immune responses^61^. Similar lymphocyte clustering has been observed with PBMCs in LN- on-chips, where flow was found to induce lymphocyte follicle formation and germinal centre characteristics without immune activation^27,28^. In our model, these immune cell clusters represent an early setting necessary for immune communication and activation. Details of lymphocyte characterisation and uncovering the mechanisms behind cluster formation, such as investigating chemoattractant-rich zones or lymphatic vessel hotspots of transendothelial migration^62,63^, could be investigated further with longer culture periods.

The LECs in the LN-on-chip model displayed similar characteristics to LN lymphatics by retaining their CD31^+^PDPN^+^ phenotype and upregulating specific LN subset markers, such as ACKR4 for SCS ceiling LECs and MARCO for medullary LECs^10^. Since ACKR4 acts as a decoy receptor for CCL21 in the SCS ceiling LECs, its action is to generate a CCL21 gradient over the floor LECs, allowing DC entry into the LN parenchyma^5,6^. The function of MARCO^+^ LECs in the LN medullary region are yet to be elucidated, but since MARCO recognises a plethora of ligands, it is hypothesised to crosstalk with macrophages for neutrophil recruitment^10^. It is unclear whether all the identified LEC subsets are present after enzymatic digestion of the LN tissue, but it was remarkable that the LECs regained LN-specific subset specifications when introduced in a LN-like microenvironmental setting. After culturing an endothelialised empty hydrogel-on-chip, in place of the LN model, most LECs lost their CD31^+^PDPN^+^ phenotype, indicating a possible mutual crosstalk between both stromal and hematopoietic cell types. In the absence of the LN model, flow alone could induce ACKR4 expression on LECs, which has been previously reported^64,65^. However, this may suggest that immune cell presence is additionally required for upregulation of MARCO, as medullary LECs are located in a region that is naturally rich in immune cells. Investigation into LEC culturing techniques may allow a future tailored approach to include specific LEC subsets into the LN-on-chip.

The choice in using the HUMIMIC Chip2 system to develop the LN-on-chip was due to the flexible ability to integrate a hydrogel-based organotypic model incorporating a customisable sacrificial printed structure^34^. In addition, the platform is versatile for use as a MOC, allowing medium to recirculate in a closed loop. This resulted in compartmentalised microenvironments, separating the circulating vasculature media and the media above the LN hydrogel. The measurements of biomarkers associated with cellular health, such as glucose, lactate and LDH, are commonly used within the OoC field^66^. Detection of these markers across the culture period indicated the viability and metabolic activity of cells within the chip. The LN-on-chip models with a LEC vessel had higher rates of LDH production, glucose consumption and lactate secretion compared to absent LEC vessels, indicating greater metabolic activity or higher concentrations with a greater cell presence. Across the seven days, all values decreased or plateaued between conditions, implying stable metabolic models.

A compartmentalised difference of soluble factors was also observed for cytokines and chemokines. The addition of the LEC vessel to the LN-on-chip model resulted in IL-7 increase in both compartments, CCL21 abundance in only the circulating vasculature compartment and more CXCL12 secretion specific to the LN model compartment. Also, both with and without the LEC vessel, CXCL13 was found exclusively in the LN model compartment. It is worth noting that with this method of extrinsic sampling, it is unknown whether such factors are present in excess or absent due to cell consumption or confinement within the hydrogel. However, advancements of new tissue engineering technologies may allow the sampling of the hydrogel microenvironment *in situ*^67^. Nonetheless, it is interesting that such signalling molecules stay local to their physiological LN- microenvironment, e.g. CCL21 in the vasculature and CXCL13 in the LN. Moreover, this becomes particularly relevant when connecting a second organotypic model to the LN model in a MOC. It would be valuable to explore the dynamic evolutions of these soluble factors over longer culture periods between the different compartments.

As such, this LN-on-chip with integrated lymphatics offers a versatile platform that can be manipulated for future experimental set-ups. One such example is the potential use of BECs instead of LECs to simulate blood vasculature. More so, this could also be used for functional experiments, including the migration of immune cells via the lymph or blood vasculature circuit to mimic incoming immune cell trafficking, as well as delivering soluble antigens or vaccines/immunotherapies to generate a local LN immune response. The current status of the LN-on-chip in terms of displaying a migratory-like LN microenvironment would indicate suitability for such experiments. This in turn will also allow the opportunity to generate a MOC to recapitulate a lymph-draining organ *in vitro*^26^. Integration of other organ models, such as skin and intestine, into a MOC with a LN model would allow the study of cross-organ communication and possible mechanisms behind immune-related disease pathophysiology. For example, immunocompetent models of healthy reconstructed human skin (RhS) are well-established within OoC systems^68–70^. Given the accurate representation of melanoma in RhS^71^, these models could be implemented into a skin-draining LN-on-chip to study tumour metastasis and anti-tumour immunity.

There are certain limitations that exist with this present study. Firstly, this method is low-throughput with scarce donor material, which would make it challenging to scale up for high-throughput experiments, which is a benchmark for the OoC and MPS field. Additionally, the one-week OoC culture duration falls short of the recommended 28-day period for translational research, based on the organisation for economic co-operation and development (OECD) guidelines for animal tests^72,73^. As earlier alluded to, the nature of hydrogel contraction is a limiting factor for this and could be overcome by using other ECM-scaffolds for the LN model. Also, some of the biological parameters are less physiological compared to native tissue. For example, restrictions with vessel printing resolutions means the 450 µm average LEC vessel width is more comparable to the diameter of small veins and arteries^74^, well-above native lymphatic measurements^75^. However, during an immune response, these diameters rapidly change with LN swelling due to the influx of immune cells^76^.

In summary, this human LN-on-chip with integrated lymphatics highlights the benefits of including LECs to generate a more physiological LN model. As such, this can be used and further advanced to recapitulate systemic human immunology.

## Author Contributions

Conceptualisation: AIM and REM; Methodology: AIM, JJ, SG, JJK and REM; Investigation: AIM and JJ; Validation: AIM and JJ; Software: AIM and JJ; Data curation: AIM and JJ; Writing-original draft preparation: AIM; Writing-review and editing: AIM, JJ, CMdW, LJWvdL, SG, JJK and REM; Supervision: CMdW, SG, JJK and REM; Funding acquisition: SG and REM; Ethical approval, sample logistical information: HPR and LJWvdL. All authors have read and agreed to the published version of the manuscript.

## Acknowledgements

The authors would like to thank and acknowledge the expert help of the Microscopy & Cytometry Core Facility (location Vrije Universiteit Amsterdam). We thank Kim Ober and Dr. Monique Verstegen (Erasmus MC, Rotterdam, The Netherlands) for sample logistics and Dr. Radjesh Bisoendial (Maasstad Hospital, Rotterdam, The Netherlands) for LEC-culturing recommendations. The presented work was funded by the European Union’s Horizon 2020 research and innovation programme under grant agreement No. 847551 ARCAID. JJ is supported by SMART Organ-on-Chip consortium, NWO-TTW Perspective Programme of the Dutch Research Council (NWO; project no. P19-03). CMdW is supported by Cancer Center Amsterdam (Grant No. CCA2019-9-57 and CCA2020-9-73) and the Dutch Cancer Foundation (KWF; Young Investigator Grant no. 2022-4 EXPL/14641). JJK is supported by the LymphChip consortium, NWA-ORC programme of the Dutch Research Council (NWO; project no. 1292.19.019). LJWvdL is supported by funding from the Convergence Health Technology Flagship grant (Organ Transplantation).

---

**Table S1.** Human lymph node donor characteristics. F: Female, M: Male, DBD: Donation after Brain Death, DCD: Donation after Circulatory Death.

| # | Donor | Sex | Age | HLA-type |
| --- | --- | --- | --- | --- |
| 1 | DBD | F | 59 | A3 A28 A68 B12 B44 B70 B72 Bw4 Bw6 Cw2 Cw5 DR3 DR17 DR4 DR52 DR53 DQ2 DQ3 DQ7 DQA-03 DQA-05 DP-0401 DPA-01 |
| 2 | DCD | F | 46 | A1 A11 B17 B57 B35 Bw4 Bw6 Cw4 Cw6 DR1 DR7 DQ1 DQ5 DQ3 DQ9 DQA-01 DQA-02 DP-0201 DPA-01 |
| 3 | DCD | M | 50 | A1 A19 A33 B14 B65 B40 B61 Bw6 Cw2 Cw8 DR3 DR17 DR6 DR13 DR52 DQ1 DQ6 DQ2 DQA-01 DQA-05 DP-0201 DP-10 DPA-01 DPA-02 |
| 4 | DBD | F | 68 | A2, A68 (28), Bw4, Bw6, B7, B53, Cw4, Cw7, DP-0401, DR52, DPA-01, DQ6 (1), DQ4, DQA-01, DQA-04, DR13 (6), DR8 |

