## Supplementary Files for "Integration of lymphatic vasculature to a human lymph node-on-chip enhances physiological immune properties"

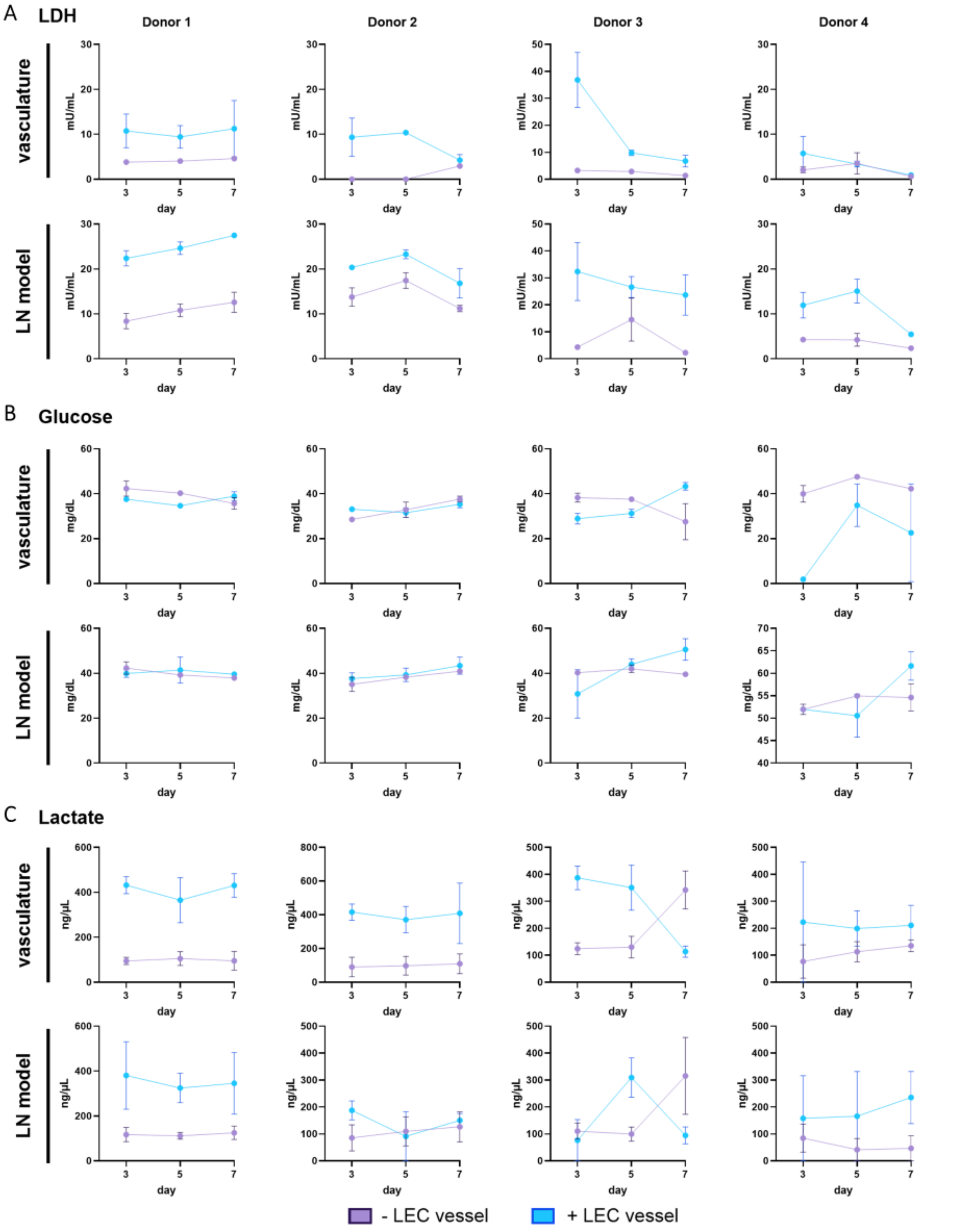


**Fig. S1: Metabolic readouts of individual donors. A.** LDH secretion **B.** Glucose concentration and **C.** Lactate secretion into culture supernatant of LN-on-chip cultures. Depicted are individual values for 4 different LN donors in duplicates.

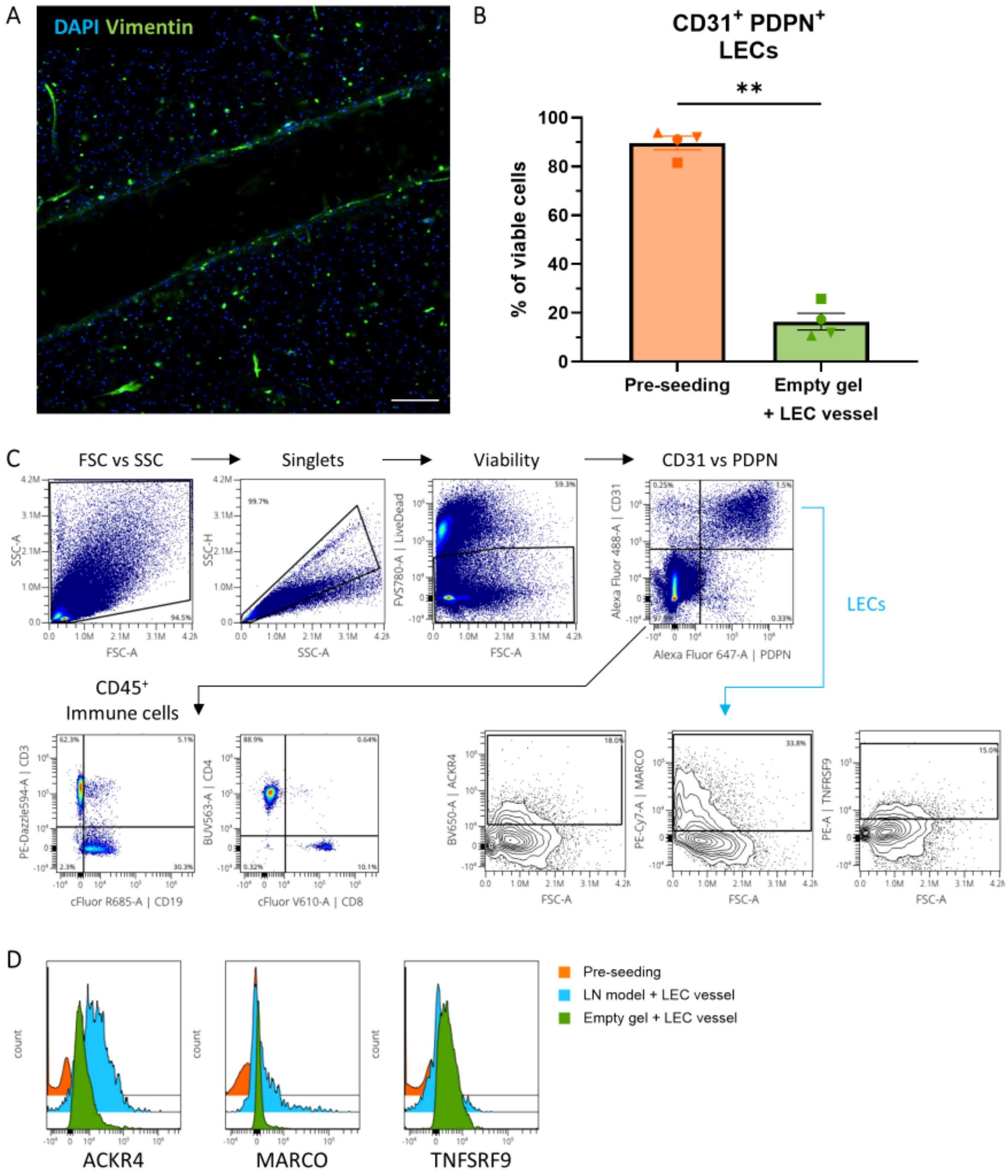


**Fig. S2: LEC characterization before seeding and in the chip. A.** Vimentin (green) and DAPI (blue) staining of a LN-on-chip with an empty vessel, imaged in the middle of the channel. Scale bar: 200 µm. **B.** LEC phenotype from CD31 and PDPN marker expression of different donors used before seeding into the chip., and after chip culture through an empty hydrogel. **C.** Representative gating strategy to identify different cells in LN model, where CD31$^{+}$PDPN$^{+}$ LECs are selected for further phenotyping (blue line), and CD31$^{-}$PDPN$^{-}$ cells are selected for immune cell gating (black line). **D.** Representative histogram overlap of LEC phenotype for ACKR4, MARCO and TNFSRF9 across pre-seeding cultured LECs, LECs in the LN model + LEC vessel, and an empty hydrogel + LEC vessel.

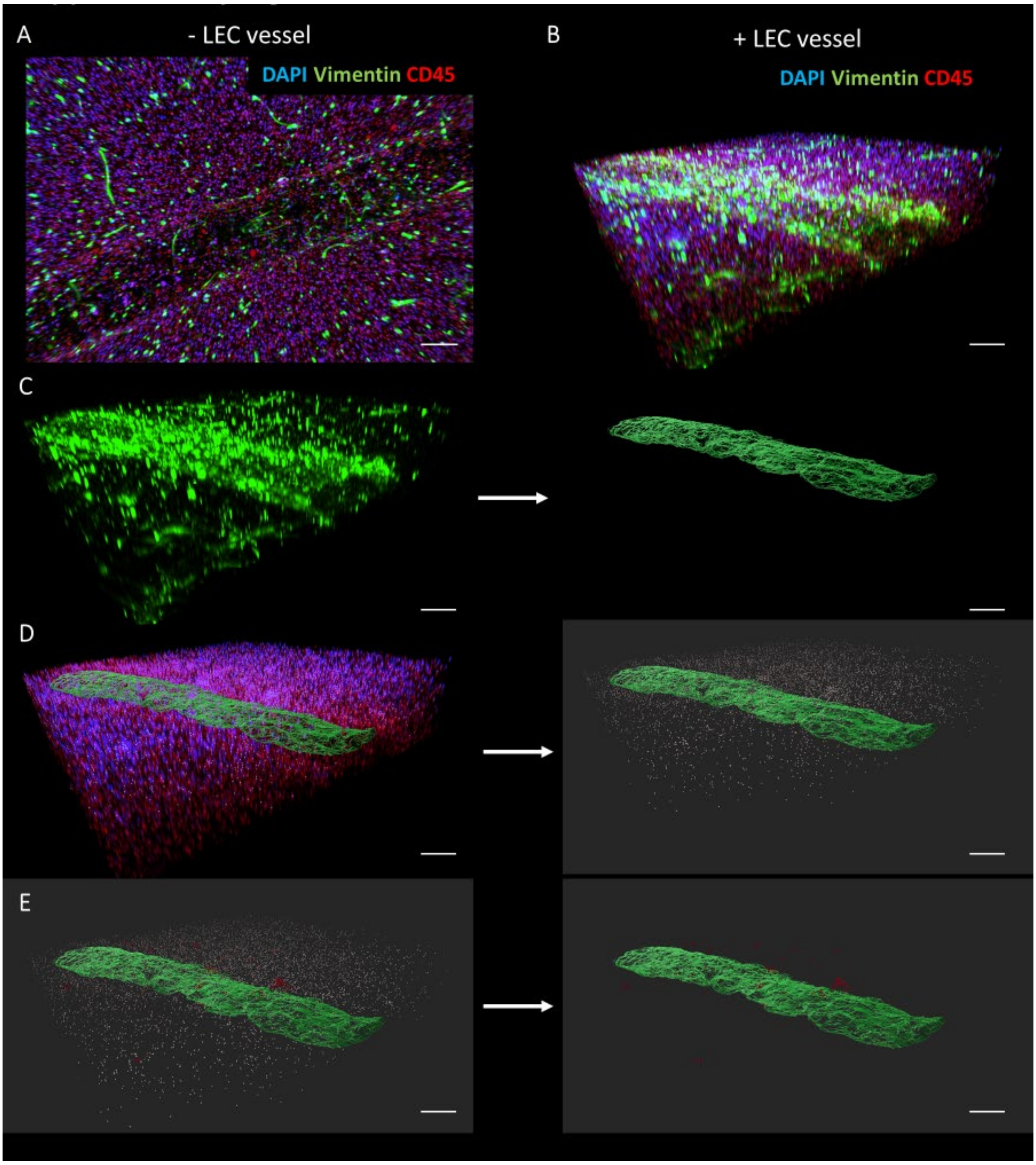


**Fig. S3: LN-on-chip renderings at day 7. A.** 3D projection of LN-on-chip without lymphatic vessel. **B.** 3D reconstruction of the LN-on-chip with lymphatic vessel stained with DAPI (blue), vimentin (green and CD45 (red)). **C.** Vessel surface rendering (green) based on vimentin. **D.** Immune cell rendering (grey) of DAPI$^+$CD45$^+$ cells. **E.** Immune cells clusters rendered (red) based on the following criteria: 9 closest neighbours within an average distance of 0-20 µm. Scale bars: 200 µm.
